# MAECI: A Pipeline For Generating Consensus Sequence With Nanopore Sequencing Long-read Assembly and Error Correction

**DOI:** 10.1101/2022.04.04.487014

**Authors:** Jidong Lang

## Abstract

Nanopore sequencing produces long reads and offers unique advantages over next-generation sequencing, especially for the assembly of draft bacterial genomes with improved completeness. However, assembly errors can occur due to data characteristics and assembly algorithms. To address these issues, we developed MAECI, a pipeline for generating consensus sequences from multiple assemblies of the same nanopore sequencing data and error correction. Systematic evaluation showed that MAECI is an efficient and effective pipeline to improve the accuracy and completeness of bacterial genome assemblies. The available codes and implementation are at https://github.com/langjidong/MAECI.

## 1. INTRODUCTION

Long reads from nanopore sequencing platforms such as Oxford Nanopore Technologies (ONT) are widely used in the study of bacterial genomes [1]. Compared with short reads from next-generation sequencing (NGS), long reads can span larger genomic repeats and complex genomic structures, thus facilitating downstream genome assembly and analysis [2, 3]. Many software or algorithm have been developed for bacterial genome assembly, such as Canu [4], FlyE [5], and Wtdbg2 [6]. They have relative advantages and disadvantages as well as varying performance and assembly outcomes [7]. Nanopore sequencing data are characterized by the presence of indels [8] and the occurrence of assembly errors spanning hundreds of bases [7], which may lead to inaccurate or incomplete assemblies. Hybrid assembly, which uses both short and long reads from next- and third-generation sequencing platforms, is gaining popularity [9]. Alternatively, the assemblies can be corrected using nanopore sequencing data [10] and then polished with NGS data. Both approaches can mitigate some of these problems and improve the accuracy of the assemblies, but assembly errors cannot be completely avoided [11]. Therefore, the assembly, especially of bacterial genomes, is far from perfect, and there are many details to consider and substantial space for improvement.

Since genome assembly is often the beginning of bioinformatics analysis by denovo sequencing of bacterial genomes, assembly errors may have critical implications for downstream analysis. Therefore, we develop MAECI, a pipeline that enables the assembly for nanopore long-read sequencing data of bacterial genomes. It takes nanopore sequencing data as input, uses multiple assembly algorithms to generate a single consensus sequence, and then uses nanopore sequencing data to perform self-error correction. MAECI takes advantage of the fact that different assembly algorithms produce different assembly errors for the same data [7], and corrects them by methods of self-correction to produce a single consensus sequence with fewer assembly error and more accurate than any of the inputs.

## 2. MATERIAL AND METHODS

### 2.1 The principles of MAECI and implementation

As shown in Figure 1, MAECI recommend that at least three assembly software and/or algorithm using long reads were used to assemble the sequencing data separately. Canu, FlyE (version: 2.8.3-b1695) and Wtdbg2 (version: 0.0 (19830203)) were selected and run with default parameters. Theoretically, the minimum number of assembly software and/or algorithm used in this step is 1, and there is no upper limit.

**Figure 1.**
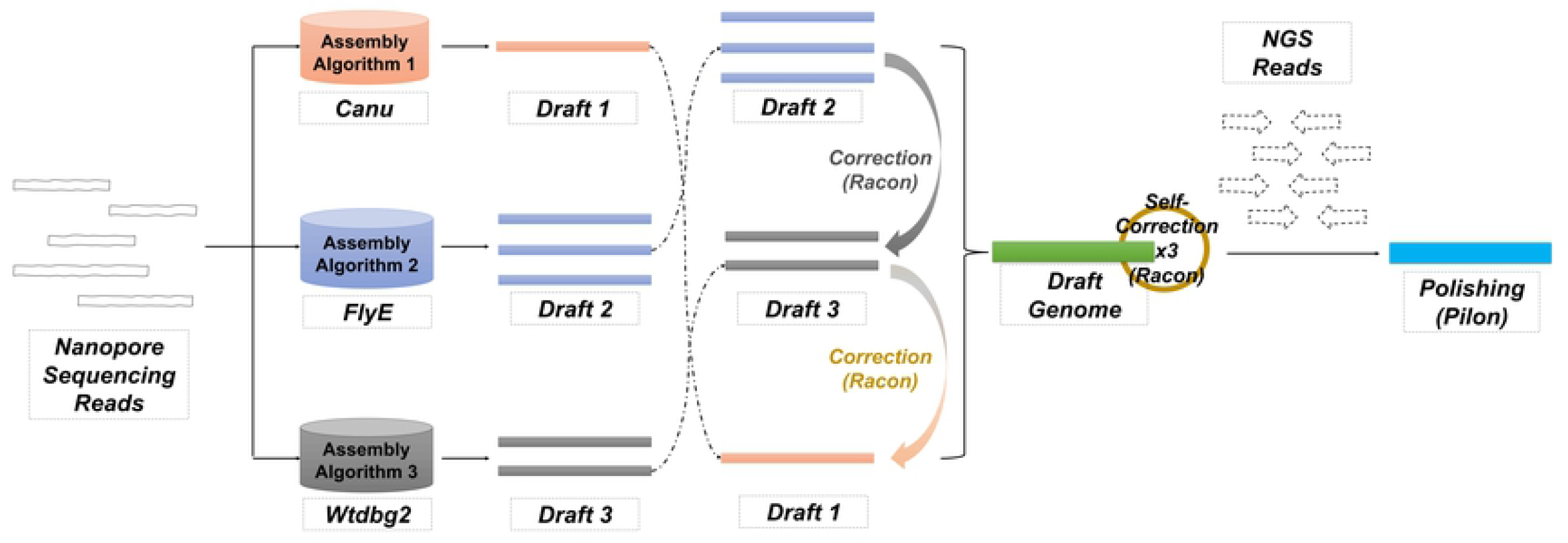
A) Overview of the MAECI assembly pipeline.

Then, the assembly results from various software and/or algorithm were sorted according to the number of scaffolds in descending order. Then, error correction algorithm, default is Racon (version: v1.4.20) [10], was used for “2+3” rounds of self-correction. For example, if the number of scaffolds from Canu, Wtdbg2, and FlyE was 3, 2, and 1, respectively, the sequencing data were aligned against the assembly from Canu in the first round of correction, with the assembly from Wtdbg2 used as the target genome to generate consensus sequence 1. In the second round of error correction, the sequencing data were aligned against consensus sequence 1, and the assembly from FlyE was used as the target genome to generate consensus sequence 2. Finally, the sequencing data and consensus sequence 2 were used for three rounds of error correction to obtain the final consensus sequence. Furtherly, if polishing with NGS data was required, default is Pilon (version: 1.24) [12], was used to polish the consensus sequence with default parameters.

By default, BWA (version: 0.7.17-r1188) [13], Minimap2 (version: 2.21-r1071) [14], Sambamba (version: 0.8.0) [15], Samtools (version: 1.12) [16] were installed for alignment. NanoPlot (version: 1.38.0) [17] was used for quality control and QUAST (version: 5.0.2) [18] was used for evaluation of the sequencing data and assemblies, with default parameters.

### 2.2 Simulated-reads tests

We downloaded nine reference bacterial strain genome sequences from NCBI: *Escherichia coli* (NC_000913.3), *Pseudomonas aeruginosa* (NC_002516.2), *Campylobacter jejuni* (NC_002163.1), *Clostridium perfringens* (NC_008261.1), *Staphylococcus aureus* (NC_007795.1), *Listeria monocytogenes* (NC_003210.1), *Enterobacter cloacae* (NZ_CP009756.1), *Faecalibacterium prausnitzii* (NZ_CP030777.1) and *Enterococcus faecalis* (NZ_KZ846041.1). Then, using NanoSim-H (version: 1.1.0.4) [19], 10,000–100,000 simulated long reads were generated from each genome (Supplementary Sheet Table S1). Using wgsim (version: 1.10) [20], simulated NGS data with a read length of 100 bp were generated for each genome (the parameters: -d 500 -s 20 -N 5000000 -e 0.01) (Supplementary Sheet Table S2).

### 2.3 Real-reads tests

We downloaded the real-world sequencing data published by *Wick et al*. [21]. The rapid sequencing and assembly results in full assemblies of three strains were selected for analysis. *trycycler_rapid_fasta*.*gz* was not found for *Serratia marcescens* 17-147-1671 dataset. For *Citrobacter koseri* MINF 9D, *Enterobacter kobei* MSB1 1B, and *Haemophilus* M1C132 1, no suitable reference genomes were available in NCBI. Thus, these three strains were excluded from analysis and comparison. As the raw ONT data were not found in the above-mentioned website, we finally downloaded the genome sequences of *Acinetobacter baumannii* strain K09-14 (NZ_CP043953.1), *Klebsiella oxytoca* strain FDAARGOS_335 (NZ_CP027426.1), and *Klebsiella variicola* strain FH-1 (NZ_CP054254.1) and used them as reference genomes. ONT data simulation was performed using NanoSim-H software. For comparison, we also performed three rounds of correction using Racon on the rapid assembly data of Trycycler [21]. QUAST was used for comparative analysis of the assemblies from MAECI and Trycycler.

## 3. RESULTS

### 3.1 Comparison of assembly results from simulated data

The contig number, total length, GC content, N50, genome fraction, mismatches per 100 kb, and indels per 100 kb of assemblies from each program were calculated using QUAST (Supplementary Sheet Table S3). MAECI was not affected by data volume, as the results were stable, and the total length and GC content after polishing using NGS data were close to the reference sequences (Supplementary Figure S1). For example, the genome of *Enterobacter cloacae* was 4,848,754 bp long with GC content of 55.03% (Figure 2A). Before polishing with NGS data, the coefficients of variation (CVs) of MAECI were 1.18e-04 (mean assembly length: 4,843,237 bp) and 2.719e-16 (mean GC content: 55.09%). After polishing, the CVs were 1.104e-05 (mean assembly length: 4,848,936 bp) and 1.361e-16 (mean GC content: 55.02%), both at the lowest. The CVs of Wtdbg2 were the highest among the four methods. Similarly, the CVs of mismatches per 100 kb of MAECI before and after polishing were 0.69 (mean: 2.945) and 0.0014 (mean: 84.522), respectively, which were the lowest. The indels per 100 kb of MAECI had CV values of 0.088 (mean: 119.668) and 0.12 (mean: 21.349) before and after polishing, respectively. MAECI and FlyE had the lowest CVs of indels per 100 kb before and after polishing, respectively. We analyzed the CVs of 10 datasets from each of the nine samples for total length, GC content, mismatches per 100 kb, and indels per 100 kb. We found that MAECI, with or without polishing with NGS data, showed low variation with different sequencing data volumes (Figure 2B).

**Figure 2.**
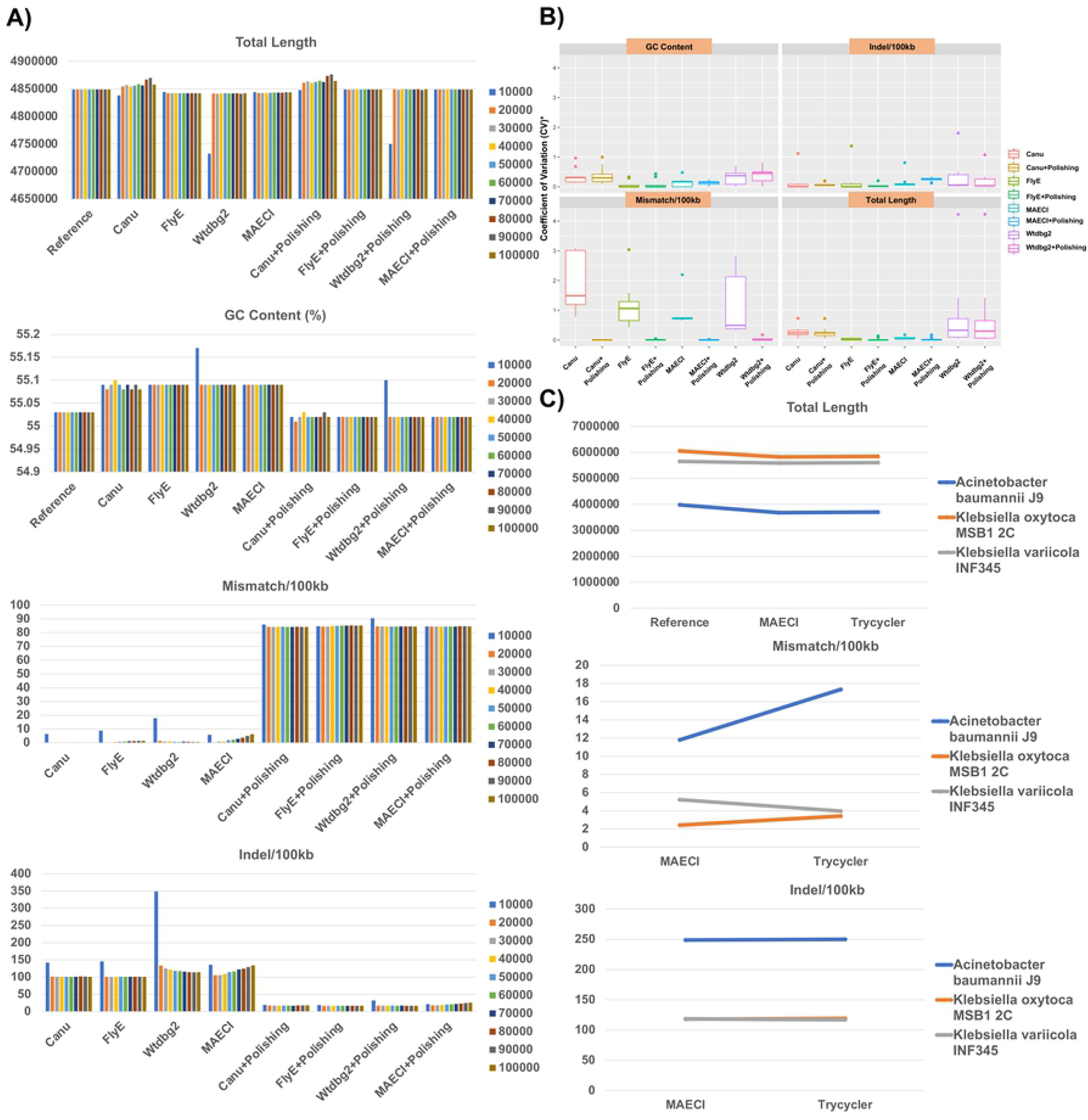
A) Comparison of the performance of Canu, FlyeE, Wtdbg2, and MAECI in the assembly of 10 simulated ONT datasets of *Enterobacter cloacae* strain GGT036 (NZ_CP009756.1) before and after polishing with simulated NGS data. B) Box plots of coefficients of variation (CVs) of total length, GC content, mismatches per 100 kb, and indels per 100 kb from 10 datasets of each of the nine samples. * represents that the smaller CVs have been enlarged: the CVs of the total length were multiplied by 100, and the GC content was multiplied by 1,000. C) Comparison of MAECI and Trycycler on real ONT datasets of three strains.

### 3.2 Comparison of assembly results from published data

The performance of MAECI was evaluated using the ONT data of *Acinetobacter baumannii* J9, *Klebsiella oxytoca* MSB1 2C, and *Klebsiella variicola* INF345 published by *Wick et al*. The number of simulated reads were close to the reported number after filtration using Filtlong (Table 1). We also performed a comparative analysis using the assemblies from Trycycler published in this article (Supplementary Sheet Table S4). We found the assembly performance of MAECI and Trycycler was largely the same, though MAECI had a lower value of mismatches per 100 kb than Trycycler, suggesting an improvement in assembly accuracy (Figure 2C, Supplementary Figure S2).

## 4. DISCUSSION

MAECI, which combined multiple input assemblies into a consensus sequence and performed self-error correction, produced a more accurate long-read-only assembly sequence. Although the results of MAECI showed the advantages of stability and high efficiency, there are still some shortcomings in the current version. In the present study, only three assembly software (Canu, FlyE and Wtdbg2) were employed, and other software and/or algorithm such as Raven [22] and La Jolla Assembler (LJA) [23] can be included for analysis, optimization and performance comparison in the next work. After polishing with NGS data, mismatches per 100 kb increased, but indels per 100 kb decreased significantly. Although this may be explained by the fact that the error rate and base or read quality score of simulated data were both “K”, the specific reasons are not further discussed in this study but will be a focus of our future research. MAECI needed more computational resources and time than single-assembler assemblies, so there may be some requirements for computing hardware.

Overall, we hope that MAECI can help to improve the accuracy of bacterial genome assembly and provide a reference solution for bacterial genome assembly. With improvements in nanopore sequencing technology, basecalling, assembly algorithms and/or polishing/error correction algorithms, it may be possible to make perfect bacterial assembly a reality.

## 5. CONCLUSIONS

MAECI is an efficient and effective pipeline for the assembly of bacterial genomes. It can be used in the assembly and analysis of bacterial and small genomes as an integration pipeline and supplementary method to genome assembly methods. With further development and optimization, we hope that this pipeline can be used in a wider range of scenarios and even in the assembly of eukaryotic or polyploid genomes.

---

**Table S1.** The simulated data statistics of three bacterial strains.

| Sample | Paper_data-Rapid<br>(after Filtlong) |  |  | Simulated data |  |  |
| --- | --- | --- | --- | --- | --- | --- |
|  | Read<br>count | Total size | N50<br>length | Read<br>count | Total size | N50<br>length |
| Acinetobacter<br>baumannii J9 | 69,135 | 741,274,205 | 15,130 | 70,000 | 543,179,080 | 9,385 |
| Klebsiella<br>oxytoca MSB1<br>2C | 69,978 | 768,514,620 | 15,181 | 70,000 | 543,767,980 | 9,411 |
| Klebsiella<br>variicola<br>INF345 | 85,440 | 948,228,758 | 15,364 | 90,000 | 698,317,203 | 9,363 |

**Figure S1.**
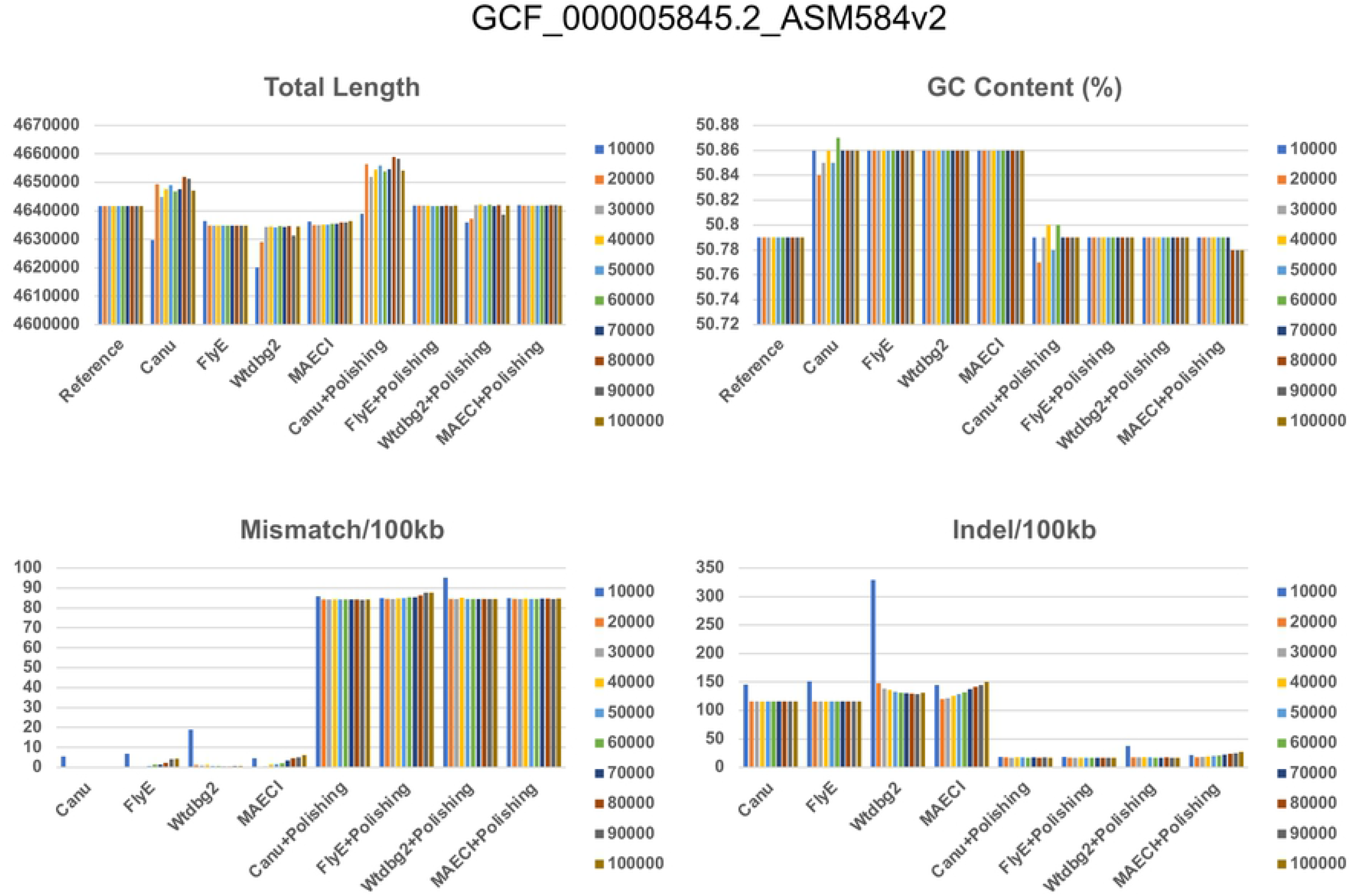
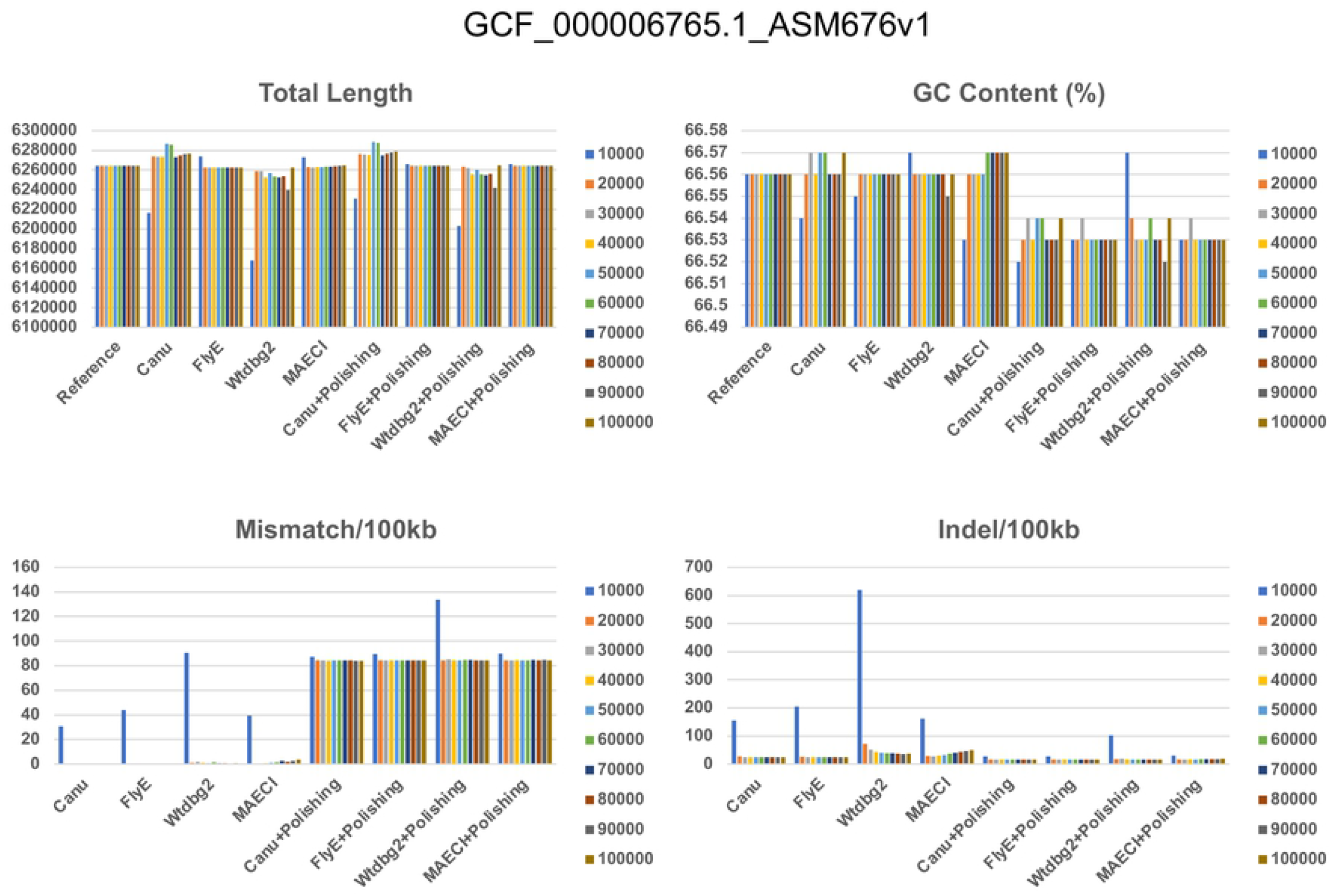
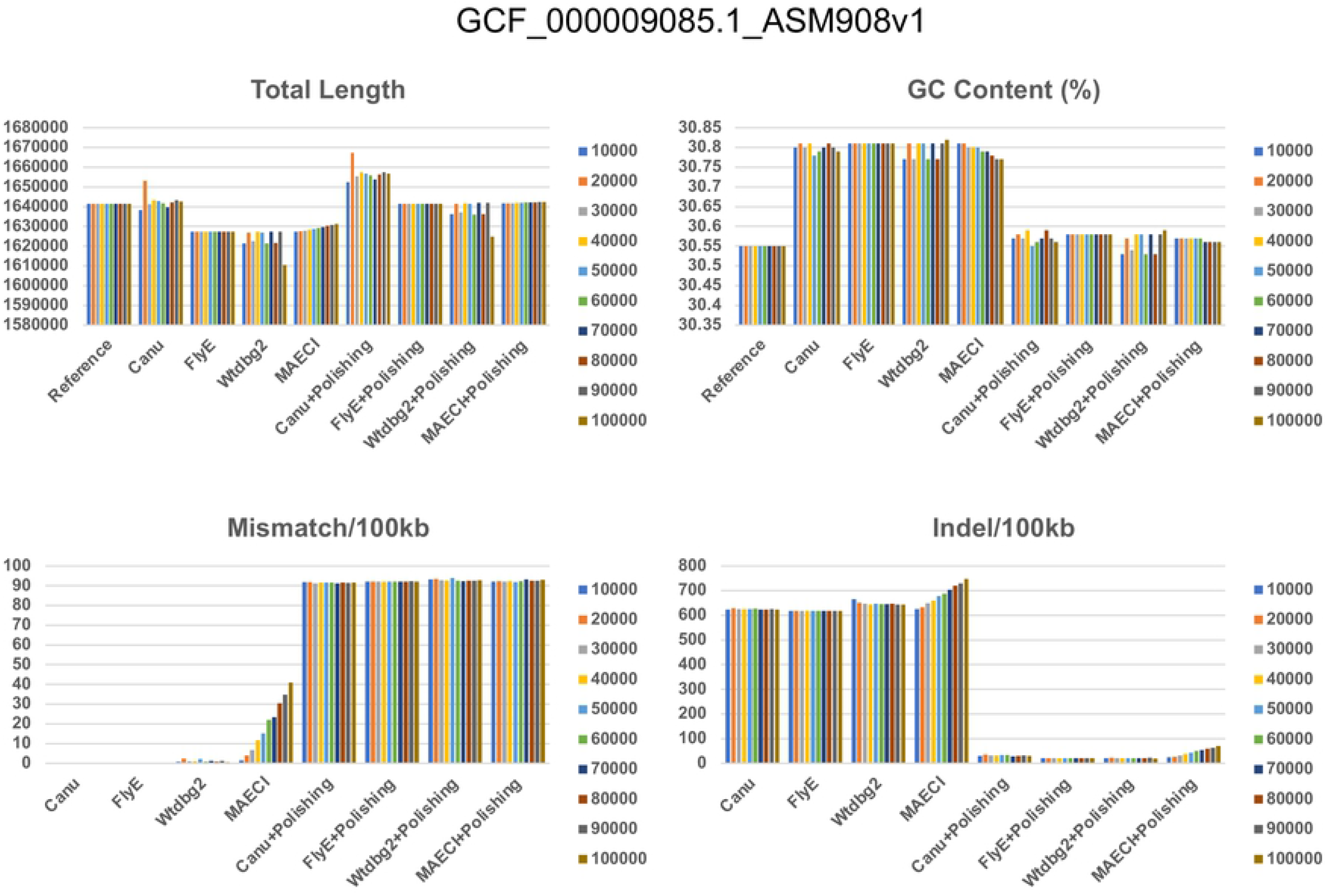
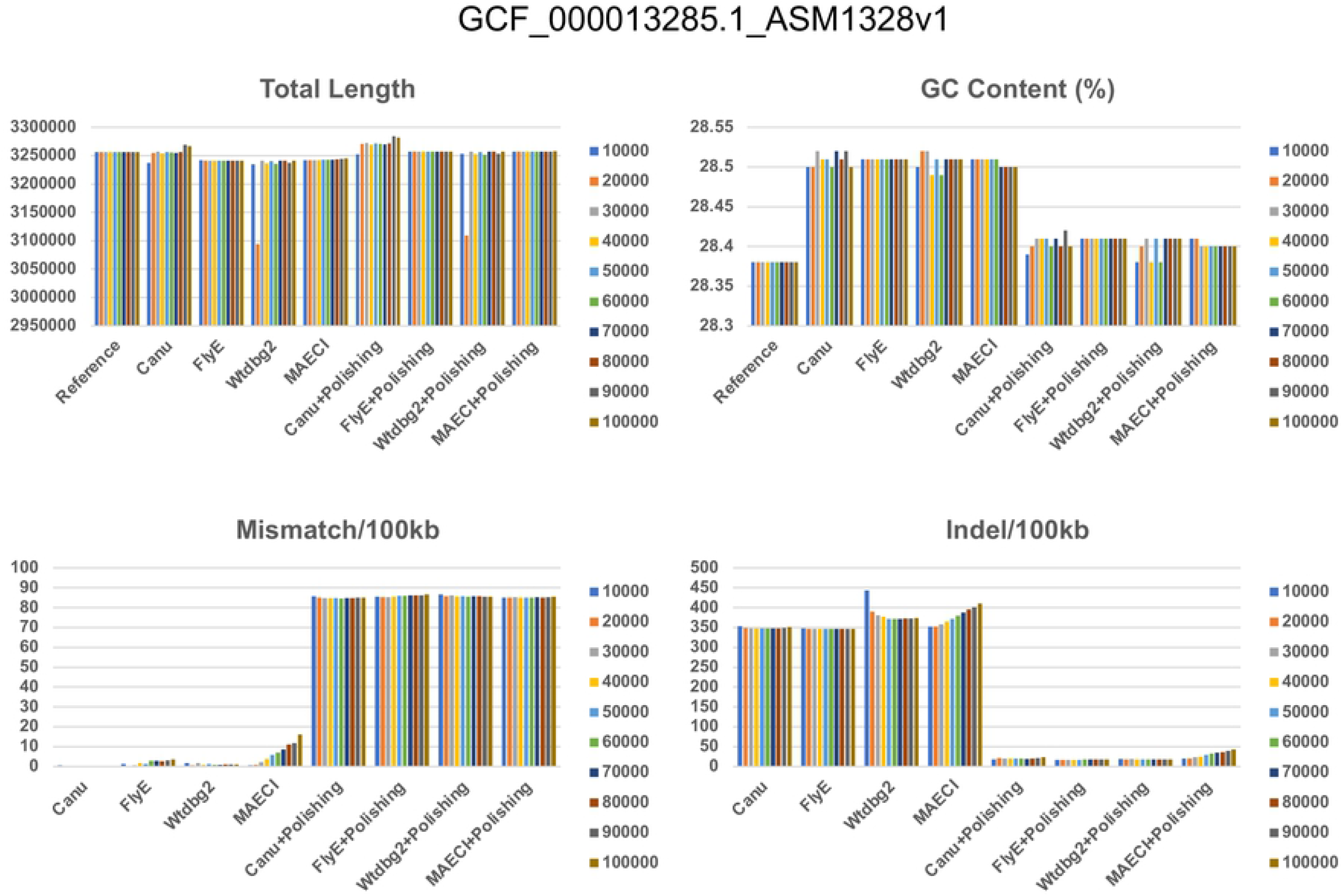
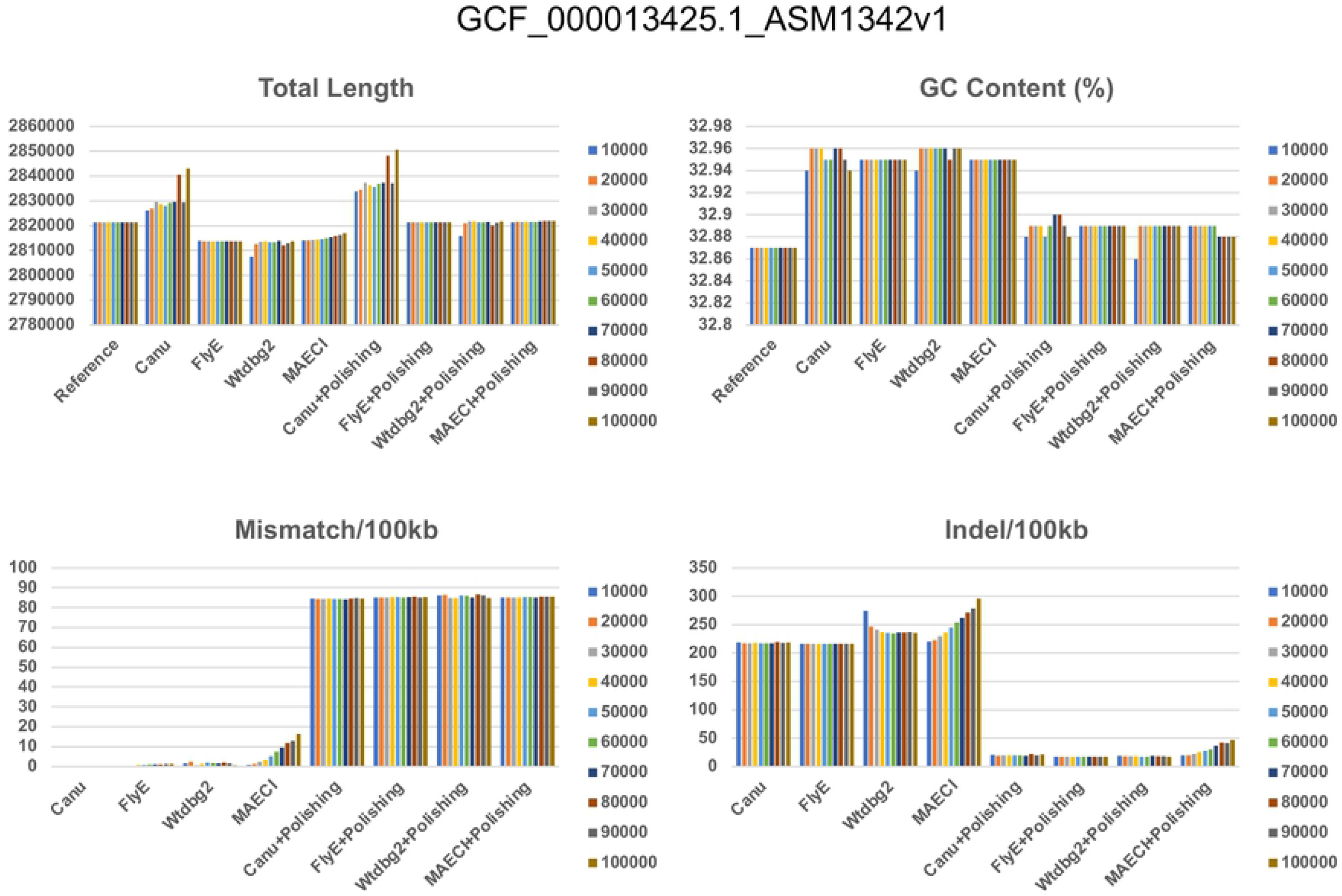
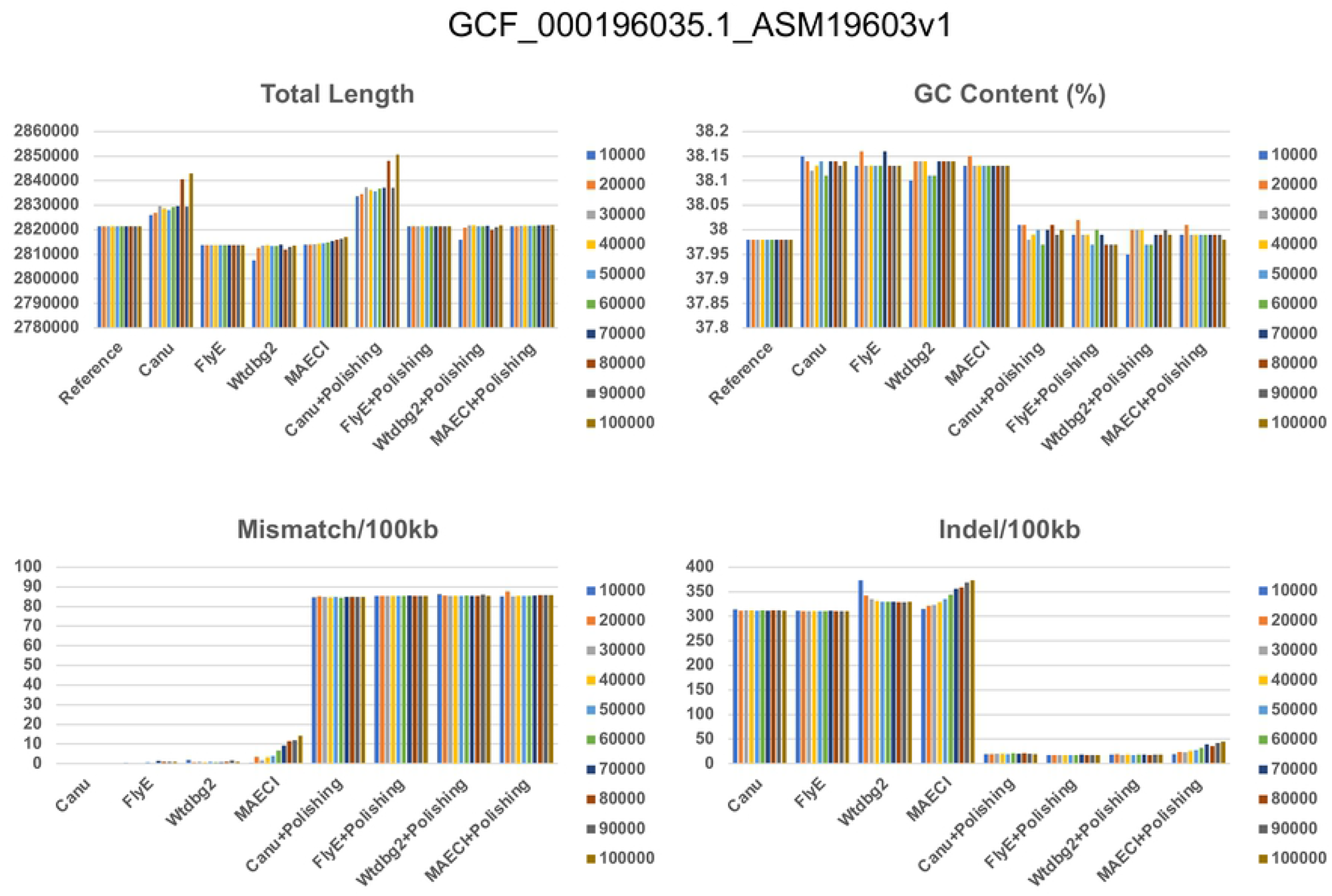
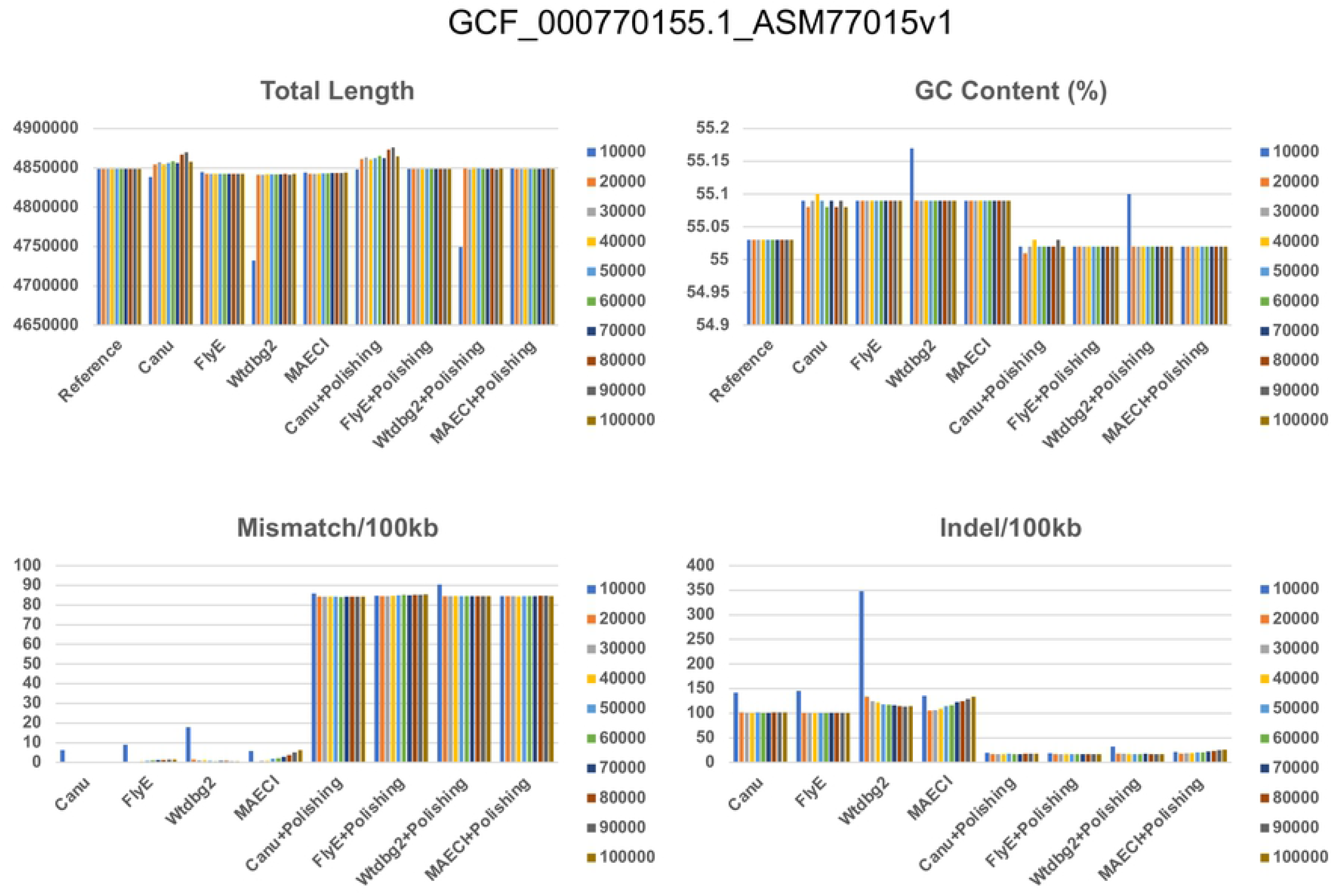
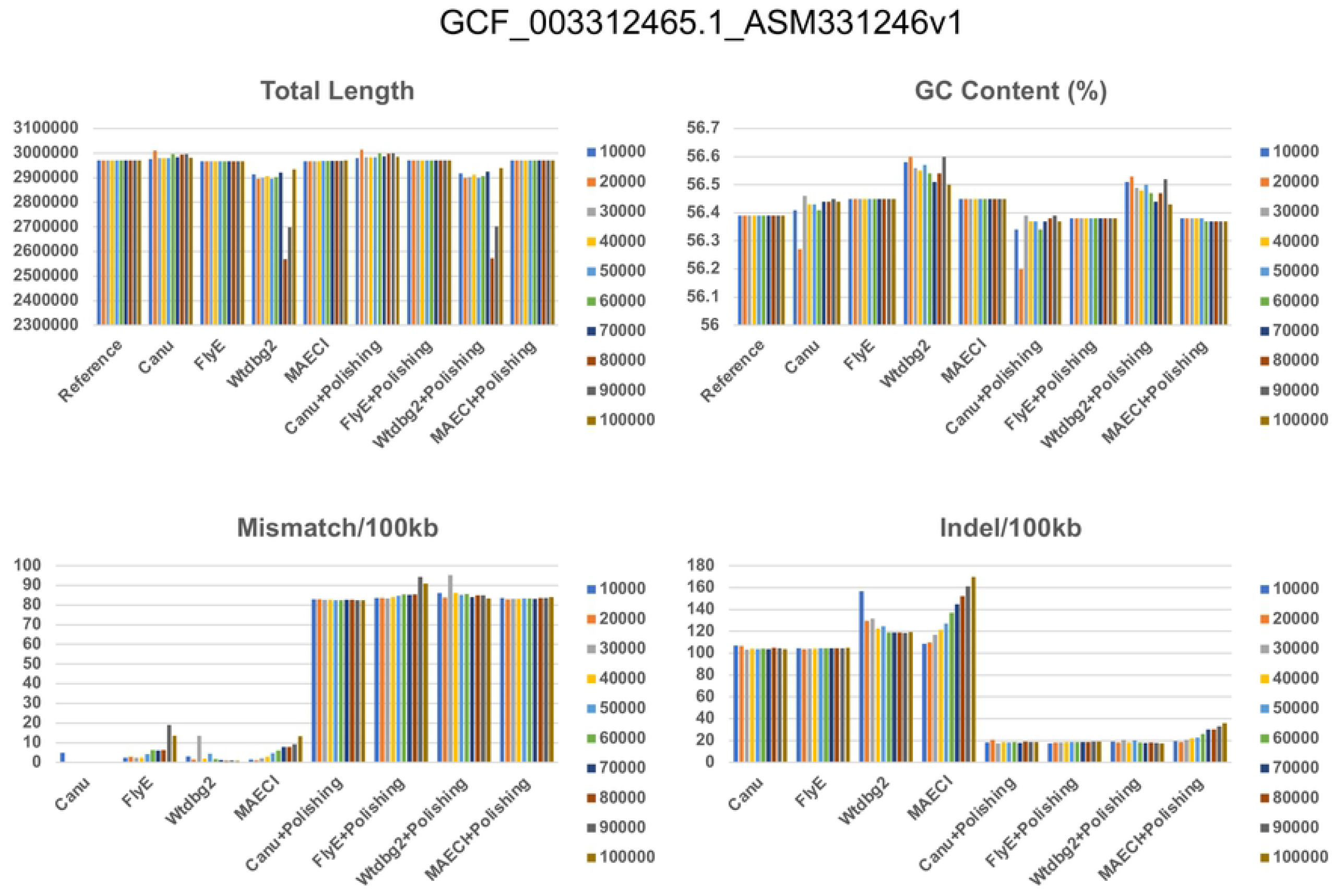
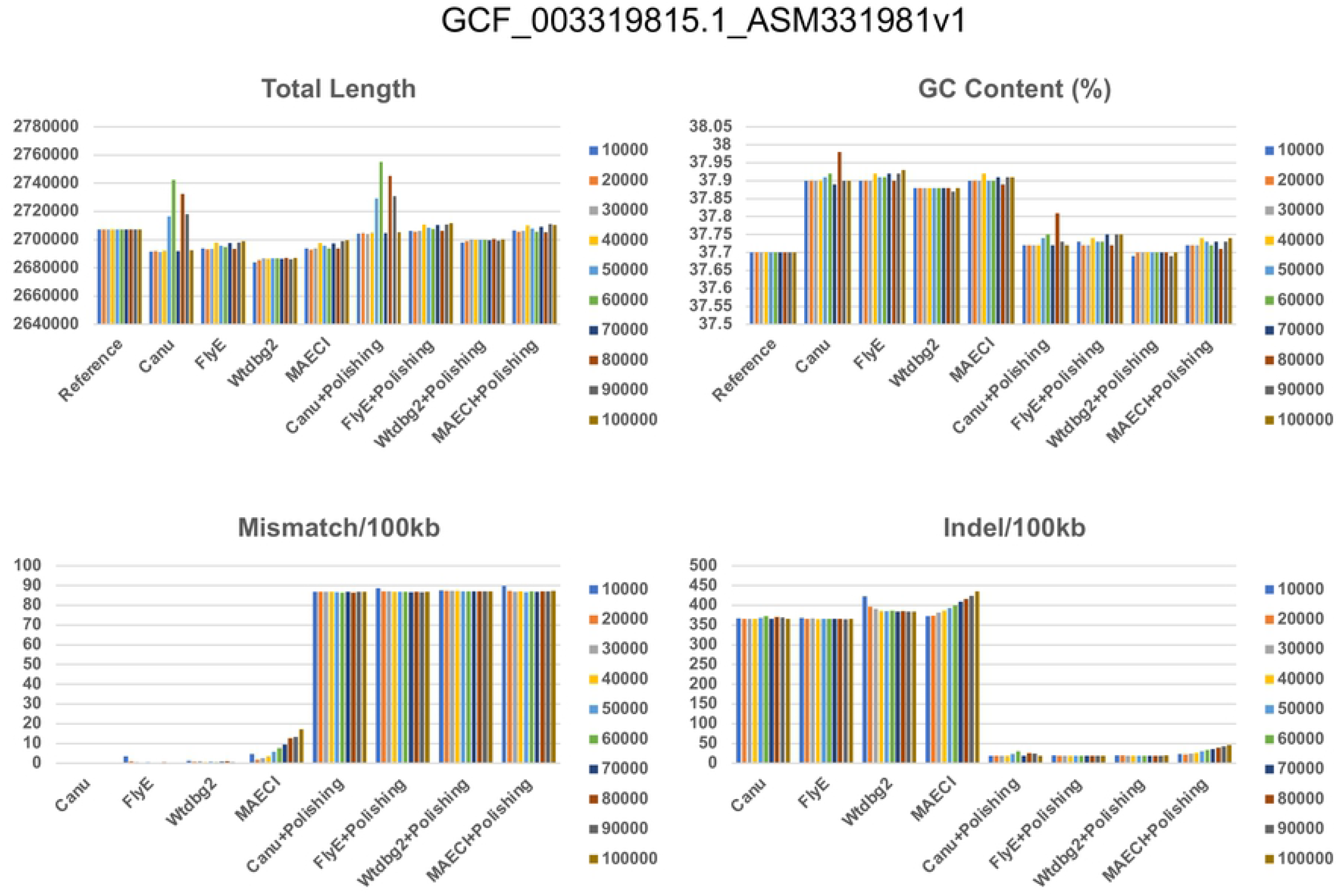
Statistics distribution chart of the bacterial strains’ assembly results.

**Figure S2.**
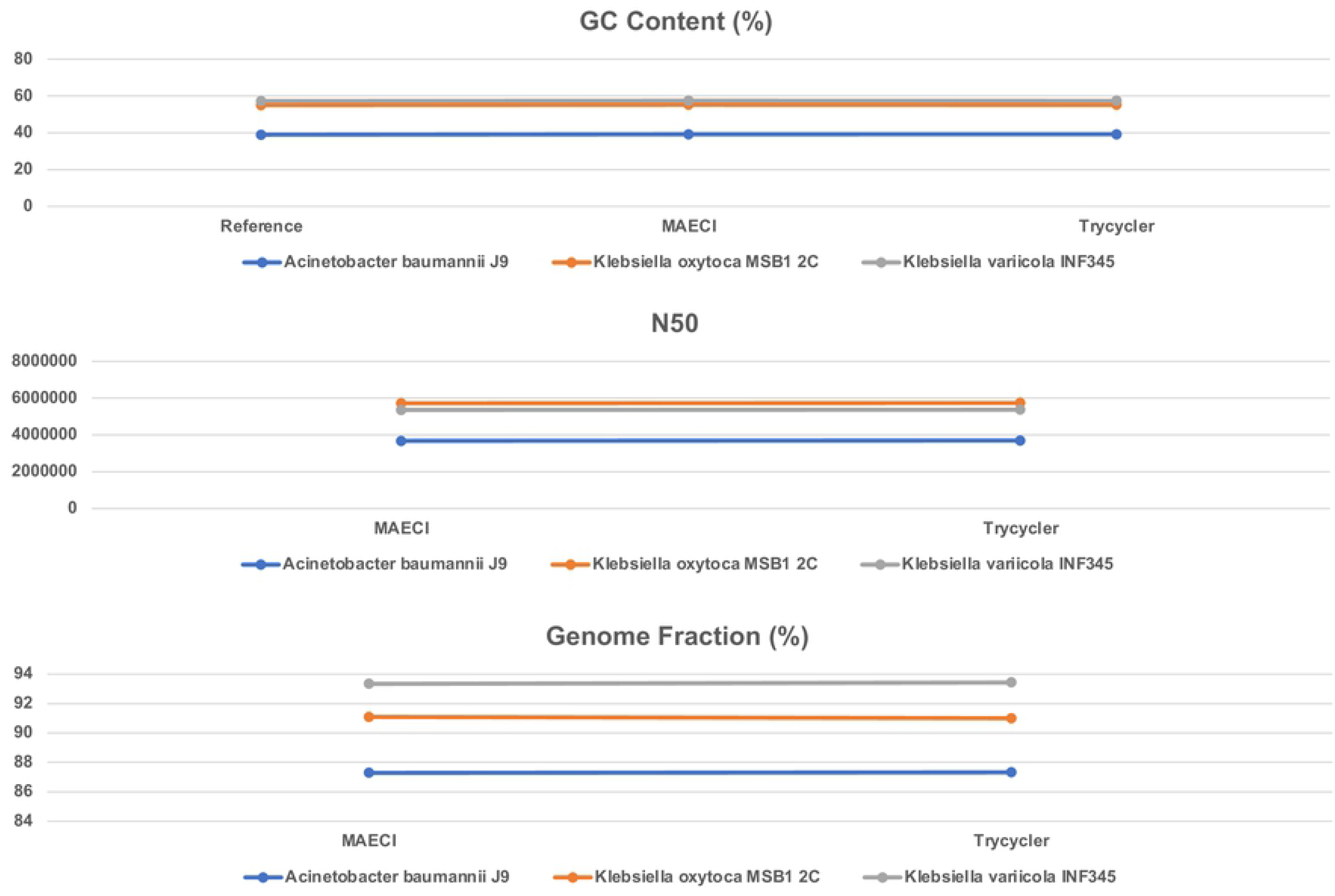
Comparison of assembly results between MAECI and Trycycler on the real data.

## REFERENCES

1. Loman, N. J., Quick, J., & Simpson, J. T. (2015). A complete bacterial genome assembled de novo using only nanopore sequencing data. Nature methods, 12(8), 733–735.

2. Miga, K. H., Koren, S., Rhie, A., Vollger, M. R., Gershman, A., Bzikadze, A., Brooks, S., Howe, E., Porubsky, D., Logsdon, G. A., Schneider, V. A., Potapova, T., Wood, J., Chow, W., Armstrong, J., Fredrickson, J., Pak, E., Tigyi, K., Kremitzki, M., Markovic, C., … Phillippy, A. M. (2020). Telomere-to-telomere assembly of a complete human X chromosome. Nature, 585(7823), 79–84.

3. Jung, H., Winefield, C., Bombarely, A., Prentis, P., & Waterhouse, P. (2019). Tools and Strategies for Long-Read Sequencing and De Novo Assembly of Plant Genomes. Trends in plant science, 24(8), 700–724.

4. Koren, S., Walenz, B. P., Berlin, K., Miller, J. R., Bergman, N. H., & Phillippy, A. M. (2017). Canu: scalable and accurate long-read assembly via adaptive k-mer weighting and repeat separation. Genome research, 27(5), 722–736.

5. Kolmogorov, M., Yuan, J., Lin, Y., & Pevzner, P. A. (2019). Assembly of long, error-prone reads using repeat graphs. Nature biotechnology, 37(5), 540–546.

6. Ruan, J., & Li, H. (2020). Fast and accurate long-read assembly with wtdbg2. Nature methods, 17(2), 155–158.

7. Wick, R. R., & Holt, K. E. (2019). Benchmarking of long-read assemblers for prokaryote whole genome sequencing. F1000Research, 8, 2138.

8. Magi, A., Giusti, B., & Tattini, L. (2017). Characterization of MinION nanopore data for resequencing analyses. Briefings in bioinformatics, 18(6), 940–953.

9. Wick, R. R., Judd, L. M., Gorrie, C. L., & Holt, K. E. (2017). Unicycler: Resolving bacterial genome assemblies from short and long sequencing reads. PLoS computational biology, 13(6), e1005595.

10. Vaser, R., Sović, I., Nagarajan, N., & Šikić, M. (2017). Fast and accurate de novo genome assembly from long uncorrected reads. Genome research, 27(5), 737–746.

11. Sović, I., Križanović, K., Skala, K., & Šikić, M. (2016). Evaluation of hybrid and non-hybrid methods for de novo assembly of nanopore reads. Bioinformatics (Oxford, England), 32(17), 2582–2589.

12. Walker, B. J., Abeel, T., Shea, T., Priest, M., Abouelliel, A., Sakthikumar, S., Cuomo, C. A., Zeng, Q., Wortman, J., Young, S. K., & Earl, A. M. (2014). Pilon: an integrated tool for comprehensive microbial variant detection and genome assembly improvement. PloS one, 9(11), e112963.

13. Li, H., & Durbin, R. (2009). Fast and accurate short read alignment with Burrows-Wheeler transform. Bioinformatics (Oxford, England), 25(14), 1754–1760.

14. Li H. (2018). Minimap2: pairwise alignment for nucleotide sequences. Bioinformatics (Oxford, England), 34(18), 3094–3100.

15. Tarasov, A., Vilella, A. J., Cuppen, E., Nijman, I. J., & Prins, P. (2015). Sambamba: fast processing of NGS alignment formats. Bioinformatics (Oxford, England), 31(12), 2032–2034.

16. Li, H., Handsaker, B., Wysoker, A., Fennell, T., Ruan, J., Homer, N., Marth, G., Abecasis, G., Durbin, R., & 1000 Genome Project Data Processing Subgroup (2009). The Sequence Alignment/Map format and SAMtools. Bioinformatics (Oxford, England), 25(16), 2078–2079.

17. De Coster, W., D’Hert, S., Schultz, D. T., Cruts, M., & Van Broeckhoven, C. (2018). NanoPack: visualizing and processing long-read sequencing data. Bioinformatics (Oxford, England), 34(15), 2666–2669.

18. Gurevich, A., Saveliev, V., Vyahhi, N., & Tesler, G. (2013). QUAST: quality assessment tool for genome assemblies. Bioinformatics (Oxford, England), 29(8), 1072–1075.

19. Yang, C., Chu, J., Warren, R. L., & Birol, I. (2017). NanoSim: nanopore sequence read simulator based on statistical characterization. GigaScience, 6(4), 1–6.

20. https://github.com/lh3/wgsim

21. Wick, R. R., Judd, L. M., Cerdeira, L. T., Hawkey, J., Méric, G., Vezina, B., Wyres, K. L., & Holt, K. E. (2021). Trycycler: consensus long-read assemblies for bacterial genomes. Genome biology, 22(1), 266.

22. Vaser, R., Šikić, M. (2021). Time- and memory-efficient genome assembly with Raven. Nature Computational Science 1, 332–336.

23. Bankevich, A., Bzikadze, A. V., Kolmogorov, M., Antipov, D., & Pevzner, P. A. (2022). Multiplex de Bruijn graphs enable genome assembly from long, high-fidelity reads. Nature biotechnology, 10.1038/s41587-022-01220-6. Advance online publication.

